# Integrating epigenomic features reveals principles of chromatin-state organization

**DOI:** 10.64898/2026.07.23.740373

**Authors:** Joseph Martini, Ryan A Williams, Rebecca G Smith, Yan Zhou, Yu Liu

## Abstract

The coordinated activities of histone modifications and chromatin-associated proteins establish chromatin states that regulate genome function and cellular identity. However, the organizational principles that distinguish chromatin states across cell types remain incompletely understood. Here, we integrated genome-wide profiles of CTCF, H3K27ac, H3K9ac, H3K27me3, and H3K9me3 at 1-kb resolution to generate a unified representation of chromatin organization in human cells. Unsupervised embedding resolved five principal chromatin states corresponding to constitutive heterochromatin, CTCF-associated architectural chromatin, transcriptionally active chromatin, mixed repressive chromatin, and Polycomb-associated chromatin. Comparative analyses of HCT116 and K562 cells revealed that cell-type-specific epigenomic differences arise predominantly through remodeling of Polycomb-associated chromatin, whereas the remaining chromatin states exhibit broadly similar epigenomic signatures and comparatively limited remodeling. Consistent with this observation, principal component analysis identified H3K27me3 as the primary contributor to genome-wide epigenomic divergence, whereas CTCF represented a secondary contributor. Integration with Hi-C data further demonstrated that CTCF-associated chromatin is strongly enriched at chromatin loop anchors and other local architectural features, linking chromatin-state organization to three-dimensional genome architecture. Together, our findings identify distinct chromatin-state classes that organize the human epigenome and reveal Polycomb-associated chromatin as a key determinant of cell-type-specific epigenomic differences.

## Introduction

Cellular identity and genome function are established through coordinated chromatin states that regulate gene expression, DNA replication, genome stability, and three-dimensional (3D) genome organization(Bannister and Kouzarides 2011; Allis and Jenuwein 2016; Bonev and Cavalli 2016). These chromatin states arise from the combinatorial activities of histone modifications and chromatin-associated proteins rather than individual epigenetic features acting in isolation. Histone acetylation marks, including H3K27ac and H3K9ac, are hallmarks of active promoters and enhancers, whereas H3K27me3 and H3K9me3 define distinct transcriptionally repressive chromatin environments associated with Polycomb-mediated repression and constitutive heterochromatin, respectively(Black et al. 2012; Consortium et al. 2020). In parallel, the architectural protein CTCF organizes higher-order chromosome structure by establishing chromatin boundaries, facilitating cohesin-mediated loop formation, and insulating regulatory interactions(Ong and Corces 2014; Nora et al. 2017; Hansen 2020). Together, these complementary chromatin features define integrated chromatin states that coordinate transcriptional regulation with higher-order genome organization.

Large-scale epigenomic mapping projects have generated comprehensive genome-wide profiles of histone modifications and chromatin-associated proteins across diverse cell types, creating unprecedented opportunities to investigate how chromatin states are established and remodeled(Roadmap Epigenomics et al. 2015; Consortium et al. 2020). Although the functions of individual chromatin features have been extensively characterized, genome regulation emerges from their coordinated interactions rather than from any single epigenetic mark. Consequently, a fundamental challenge in epigenomics is to identify the major chromatin-state classes that organize the genome and determine which chromatin features drive cell-type-specific epigenomic organization.

Integrative computational approaches, including ChromHMM and Segway, have substantially advanced genome annotation by integrating multiple epigenomic datasets to classify promoters, enhancers, transcribed regions, and repressive chromatin domains(Ernst and Kellis 2012; Hoffman et al. 2012). These methods excel at functional genome annotation but are not intended to reveal the principal chromatin features that distinguish one epigenome from another. As increasingly diverse epigenomic datasets become available, complementary analytical strategies that integrate multiple chromatin features while preserving relationships among genomic regions may provide new insights into the organizational principles of the epigenome.

Here, we addressed this challenge by integrating genome-wide profiles of CTCF, H3K27ac, H3K9ac, H3K27me3, and H3K9me3 into a unified chromatin-state representation at 1-kb resolution. Rather than assigning genomic regions to predefined functional categories, we applied unsupervised manifold learning to identify chromatin states directly from shared epigenomic similarity(Becht et al. 2018; McInnes 2018), enabling unbiased comparisons of chromatin organization across cell types. This analysis identified five major chromatin-state classes corresponding to constitutive heterochromatin, CTCF-associated architectural chromatin, transcriptionally active chromatin, mixed repressive chromatin, and Polycomb-associated chromatin. Comparative analyses further revealed that cell-type-specific epigenomic differences arise predominantly through remodeling of Polycomb-associated chromatin, whereas CTCF-associated chromatin defines an architectural state enriched at chromatin loop anchors. Together, these findings reveal complementary principles of chromatin-state organization in which architectural chromatin provides a structural framework for genome organization, while selective remodeling of Polycomb-associated chromatin underlies cell-type-specific epigenomic diversity.

## Results

### Integrative epigenomic analysis identifies five major chromatin-state classes

To investigate how multiple chromatin features collectively organize the human epigenome, we integrated genome-wide ChIP-seq profiles of the architectural protein CTCF together with active histone modifications (H3K27ac and H3K9ac) and repressive histone modifications (H3K27me3 and H3K9me3) from HCT116 and K562 cells (Fig. 1A). These five features capture complementary aspects of chromatin architecture, transcriptional activity, facultative repression, and constitutive heterochromatin(Bannister and Kouzarides 2011; Ong and Corces 2014; Allis and Jenuwein 2016), providing a multidimensional representation of chromatin organization.

**Figure 1.**
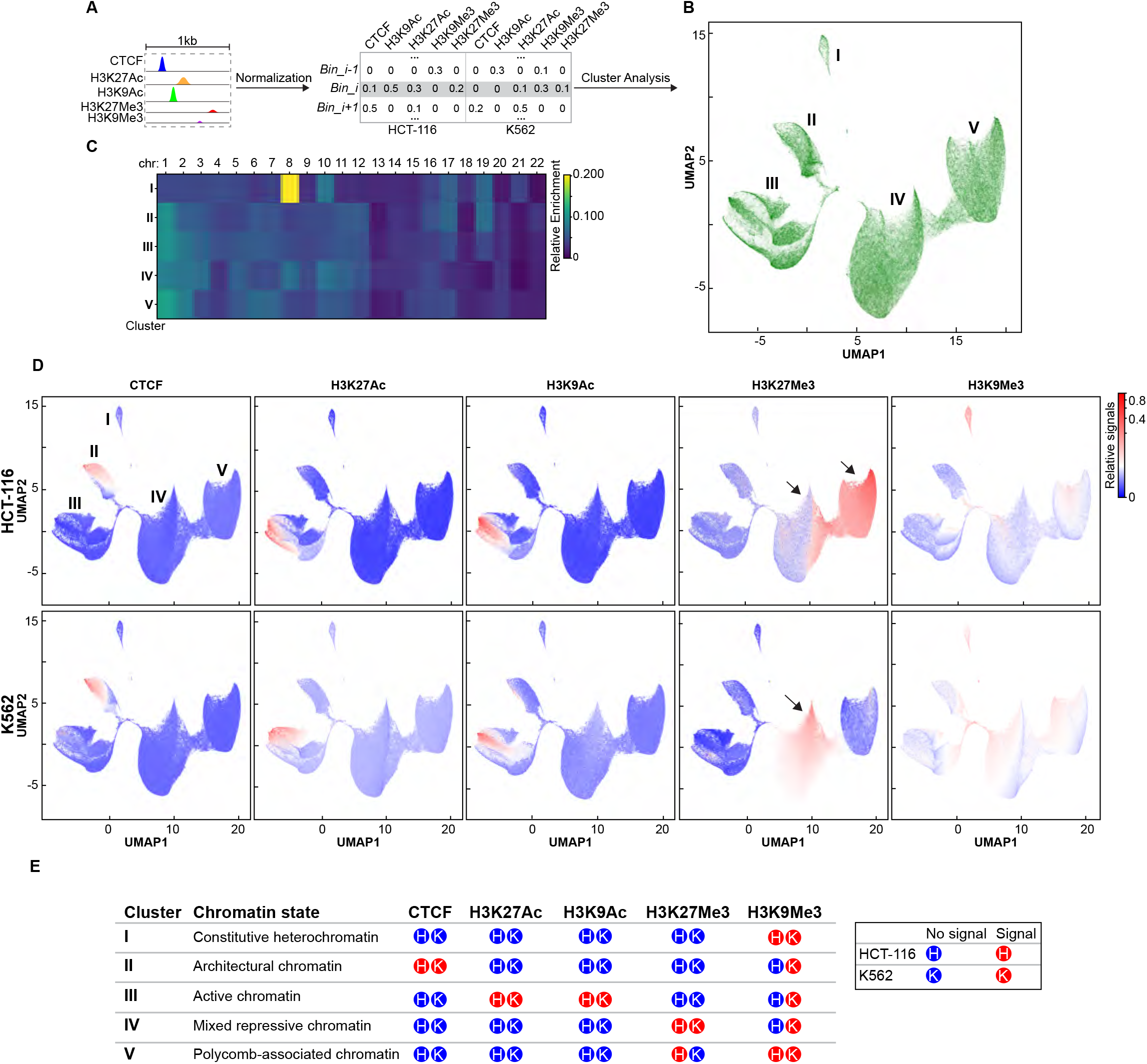
Integration of multiple chromatin features identifies distinct genome-wide chromatin states. (A)Schematic overview of the analytical workflow. The human genome was partitioned into 1-kb bins, and ChIP-seq signals for CTCF, H3K27me3, H3K9me3, H3K27ac, and H3K9ac were quantified in each bin using datasets from HCT116 and K562 cells. Following quality control and Z-score normalization, each genomic bin was represented by a five-dimensional epigenomic feature vector and analyzed using dimensionality-reduction methods commonly applied in single-cell studies. (B)UMAP projection of genomic bins based on integrated CTCF and histone modification profiles from HCT116 and K562 cells, revealing five distinct epigenomic clusters. (C)Chromosomal enrichment of genomic bins in each epigenomic cluster. (D)Projection of CTCF and histone modification signals onto the UMAP manifold. Color intensity represents normalized ChIP-seq signal for each chromatin feature. (E)Epigenomic characteristics and biological interpretation of the five chromatin clusters.

The genome was partitioned into 1-kb intervals, and each interval was represented by its integrated chromatin signature after normalization of the five epigenomic features (Methods). We then performed unsupervised manifold learning using UMAP followed by graph-based clustering to determine whether recurrent chromatin configurations emerge directly from integrated epigenomic information(Becht et al. 2018; McInnes 2018). This analysis resolved five well-defined chromatin-state classes shared by both cell types (Fig. 1B), demonstrating that combinations of chromatin features naturally partition the genome into a limited number of reproducible chromatin states.

Chromosome-level analysis revealed broadly similar genomic distributions for four of the five chromatin states. In contrast, the constitutive heterochromatin state exhibited pronounced enrichment on chromosome 8 (Fig. 1C), indicating that chromosome-specific heterochromatin contributes uniquely to this chromatin-state class.

To determine the biological identity of the five chromatin states, we examined the distribution of individual chromatin features within each cluster (Fig. 1D). Each state displayed a distinct and biologically interpretable chromatin signature.

The first chromatin state was characterized by strong H3K9me3 enrichment with minimal CTCF, H3K27ac, H3K9ac, or H3K27me3 signal in both cell types, consistent with constitutive heterochromatin(Peters et al. 2003; Saksouk et al. 2015) (Fig. 1D). The second state was distinguished by prominent CTCF enrichment, consistent with an architectural chromatin state(Ong and Corces 2014; Hansen 2020). Although CTCF occupancy was evident in both HCT116 and K562 cells, its genomic distribution differed substantially between cell types, indicating that the architectural chromatin state occupies distinct genomic location in differenct cell types (Fig. 1D).

The third chromatin state exhibited coordinated enrichment of H3K27ac and H3K9ac, consistent with transcriptionally active chromatin(Creyghton et al. 2010; Bannister and Kouzarides 2011). Active histone acetylation was largely mutually exclusive with H3K9me3, reflecting the separation of active and constitutively repressed chromatin environments. Although the same active chromatin state was identified in both HCT116 and K562 cells, the genomic locations of acetylated regions differed substantially between the two cell types.

The remaining two chromatin states both displayed repressive chromatin signatures but differed markedly in composition. One state exhibited mixed H3K27me3 and H3K9me3 signals, with substantially greater H3K9me3 enrichment in K562 cells, suggesting differential integration of constitutive and facultative repression. The other was dominated by H3K27me3 with relatively weak H3K9me3, consistent with Polycomb-associated chromatin(Margueron and Reinberg 2011; Blackledge and Klose 2021). Notably, H3K27me3 enrichment within this state was considerably stronger in HCT116 than in K562 cells, indicating extensive remodeling of Polycomb-associated chromatin between the two cell types.

Together, these analyses demonstrate that the human epigenome can be partitioned into five major chromatin-state classes representing constitutive heterochromatin, architectural chromatin, active chromatin, mixed repressive chromatin, and Polycomb-associated chromatin. While all five chromatin states are present in both cell types, Polycomb-associated chromatin exhibits the most pronounced cell-type-specific remodeling.

### Polycomb-associated chromatin is the primary source of cell-type-specific epigenomic divergence

The chromatin-state analysis suggested that Polycomb-associated chromatin, a hallmark of facultative repression(Margueron and Reinberg 2011; Blackledge and Klose 2021), contributes disproportionately to the differences between HCT116 and K562 cells. Among the five chromatin states, the Polycomb-associated state exhibited the most prominent cell-type-specific differences, whereas constitutive heterochromatin, architectural chromatin, and active chromatin displayed comparatively limited chromatin-state remodeling. These observations suggested that remodeling of Polycomb-associated chromatin represents the major source of epigenomic divergence.

To independently evaluate the contribution of individual chromatin features, we performed principal component analysis (PCA) separately for each epigenomic feature using genome-wide ChIP-seq profiles from HCT116 and K562 cells (Fig. 2A)(Jolliffe and Cadima 2016). H3K27me3 produced the greatest separation between the two cell types, indicating that Polycomb-associated chromatin differs more extensively than any other chromatin feature. Quantification of Euclidean distances within PCA space confirmed H3K27me3 as the largest contributor to genome-wide epigenomic divergence (Fig. 2B). CTCF displayed the second-largest separation, suggesting that differences in chromatin architecture also contribute to cell-type-specific chromatin organization, albeit to a lesser extent.

**Figure 2.**
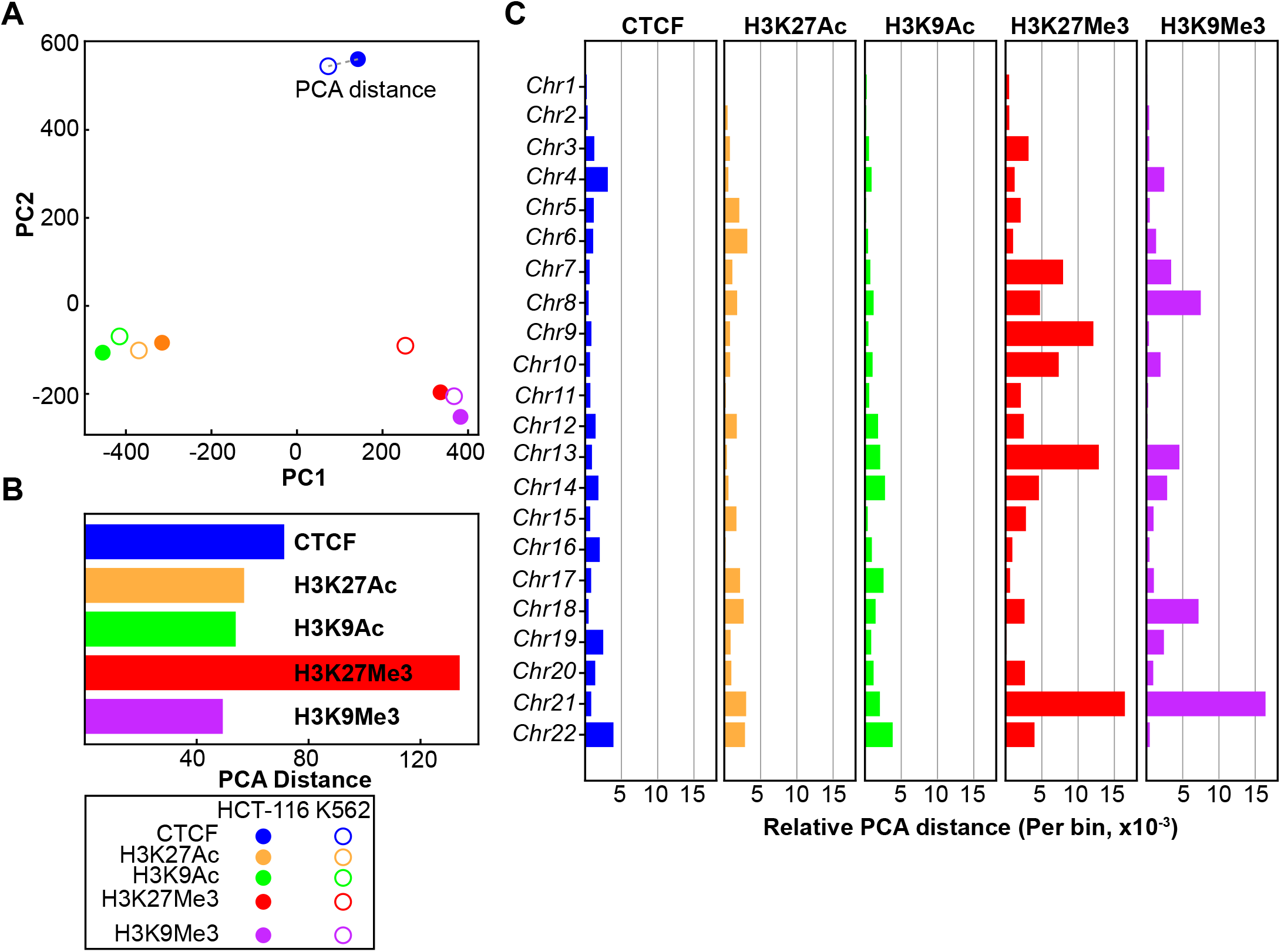
Principal component analysis identifies H3K27me3 as the dominant source of epigenomic variation between HCT116 and K562 cells. (A)Principal component analysis (PCA) of genome-wide epigenomic profiles. Solid and open circles represent genomic bins from HCT116 and K562 cells, respectively. Colors indicate the five epigenomic clusters identified by UMAP. (B)Quantification of PCA-based separation between HCT116 and K562 for individual chromatin features and CTCF. Separation was measured as the Euclidean distance between cell-type centroids in principal component space. (C)Chromosome-level PCA showing the Euclidean distance between HCT116 and K562 centroids for each chromatin feature and CTCF.

We next asked whether these differences occurred globally or were restricted to individual chromosomes. Chromosome-level PCA demonstrated that H3K27me3 consistently exhibited the greatest separation between HCT116 and K562 across most chromosomes (Fig. 2C). Particularly strong divergence was observed on chromosomes 7, 9, 13, and 21. H3K9me3 also showed chromosome-specific differences, especially on chromosomes 8, 18, and 21, but these changes were comparatively localized and did not account for the global separation observed between the two cell types.

Together, these independent analyses demonstrate that remodeling of Polycomb-associated chromatin constitutes the dominant source of epigenomic variation between HCT116 and K562 cells, whereas changes in CTCF occupancy represent a secondary axis of divergence.

### CTCF-associated chromatin defines local architectural domains

Because one chromatin-state class was defined primarily by CTCF enrichment, a key architectural protein involved in chromatin looping and genome organization(Ong and Corces 2014; Hansen 2020), we next investigated its relationship with 3D genome architecture using Hi-C data(Lieberman-Aiden et al. 2009; Dixon et al. 2012; Rao et al. 2014). The different spatial resolutions of ChIP-seq and Hi-C present a challenge for direct integration. ChIP-seq signals were analyzed at 1-kb resolution, whereas chromatin loops and TAD boundaries are typically identified at approximately 20-kb resolution and A/B compartments at even larger genomic scales. To evaluate the compatibility of these datasets, we first examined the continuity of chromatin-state assignments by quantifying genomic regions containing consecutive 1-kb bins belonging to the same chromatin state.

As expected, the frequency of continuous chromatin-state regions decreased as the required number of consecutive bins increased (Fig. 3A-C). Fewer than 10% of regions contained 20 consecutive bins, indicating that most chromatin states represent relatively local genomic features. Consequently, subsequent analyses focused on local architectural elements, including chromatin loop anchors and TAD boundaries.

**Figure 3.**
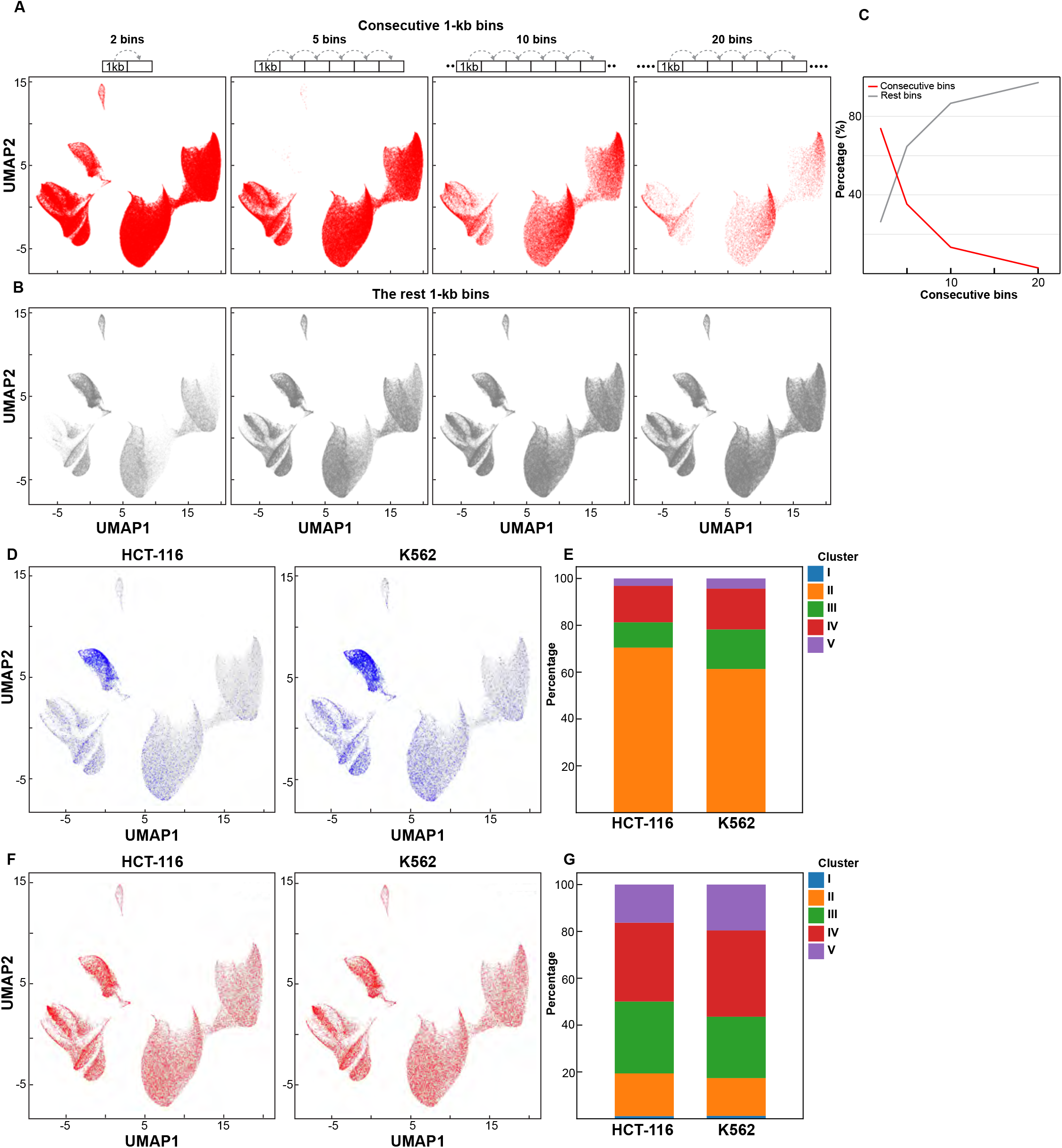
Association of epigenomic clusters with local chromatin architectural features. (A, B) Distribution of consecutive 1-kb genomic bins within each epigenomic cluster. Genomic bins belonging to runs of 2, 5, 10, or 20 consecutive bins are highlighted in red, whereas all remaining bins are shown in gray. (C) Percentage of genomic bins belonging to runs of 2, 5, 10, or 20 consecutive bins (red) and the corresponding non-consecutive bins (gray). (D, E) Enrichment of CTCF–CTCF loop-anchor bins across epigenomic clusters in HCT116 and K562 cells. (F, G) Enrichment of TAD-boundary bins across epigenomic clusters in HCT116 and K562 cells.

Integration with Hi-C data revealed a striking enrichment of CTCF-mediated loop anchors within the architectural chromatin state in both HCT116 and K562 cells (Fig. 3D and 3E), providing independent validation that this chromatin state corresponds to local architectural elements of the genome(Ong and Corces 2014; Hansen 2020). In K562 cells, active chromatin also showed moderate enrichment at loop anchors, suggesting a stronger association between chromatin looping and transcriptionally active regions in this cell type.

In contrast, TAD boundaries exhibited relatively modest enrichment across multiple chromatin states, with only a slight preference for the mixed repressive chromatin state. Overall enrichment patterns were highly similar between HCT116 and K562 cells, suggesting that TAD boundaries are less specifically associated with individual chromatin states than chromatin loop anchors (Fig. 3F and 3G). We note that the comparatively coarse resolution of TAD annotations may also reduce the sensitivity of these analyses.

Collectively, these findings demonstrate that the CTCF-associated chromatin state defines local architectural domains of the genome. Together with the identification of Polycomb-associated chromatin as the principal source of epigenomic divergence, these results reveal complementary organizational principles of the human epigenome, in which architectural chromatin provides a structural framework while selective remodeling of Polycomb-associated chromatin underlies cell-type-specific epigenomic diversity.

## Discussion

Chromatin states emerge from the coordinated actions of histone modifications and chromatin-associated proteins to regulate gene expression, genome stability, and higher-order chromosome organization(Bannister and Kouzarides 2011; Allis and Jenuwein 2016; Bonev and Cavalli 2016). Although genome-wide epigenomic maps have become increasingly comprehensive, how multiple chromatin features collectively organize the epigenome and distinguish one cell type from another has remained incompletely understood. By integrating five complementary epigenomic features into a unified chromatin-state representation, our study identifies a limited number of shared chromatin-state classes that capture the major functional and architectural domains of the human genome. More importantly, our analyses reveal that cell-type-specific epigenomic variation is not distributed uniformly across chromatin states but arises predominantly through selective remodeling of Polycomb-associated chromatin, whereas CTCF-associated chromatin forms a stable architectural scaffold linked to three-dimensional genome organization(Ong and Corces 2014; Hansen 2020).

One of the central findings of this study is that Polycomb-associated chromatin represents the primary source of epigenomic divergence between HCT116 and K562 cells. Both unsupervised chromatin-state analysis and independent principal component analysis consistently identified H3K27me3 as the dominant contributor to genome-wide differences between the two cell types. This observation is consistent with the established role of Polycomb-mediated repression in developmental gene regulation and maintenance of cellular identity(Margueron and Reinberg 2011; Blackledge and Klose 2021). Rather than being confined to a limited set of genomic loci, H3K27me3 differences were broadly distributed across chromosomes, indicating that remodeling of Polycomb-associated chromatin represents a global organizational feature of cell-type-specific epigenomes. In contrast, active chromatin, constitutive heterochromatin, and CTCF-associated chromatin exhibited comparatively limited chromatin-state remodeling, suggesting that cell-type-specific epigenomic differences are concentrated within specific chromatin states rather than distributed uniformly across the epigenome.

Our analyses also demonstrate that CTCF-associated chromatin defines an architectural chromatin state. Although chromatin states were inferred exclusively from epigenomic profiles, the CTCF-enriched state was strongly associated with chromatin loop anchors identified by Hi-C(Rao et al. 2014), providing independent validation that integrated chromatin signatures capture biologically meaningful features of 3D genome organization. In contrast, TAD boundaries exhibited comparatively weak state-specific enrichment, suggesting that their establishment depends on additional architectural determinants beyond the chromatin features analyzed here. Together, these findings indicate that integrated chromatin states provide a direct connection between one-dimensional epigenomic profiles and local chromosome architecture(Ong and Corces 2014; Hansen 2020) and support a model in which CTCF-associated architectural chromatin provides a relatively stable structural framework, whereas Polycomb-associated chromatin undergoes extensive cell-type-specific remodeling.

An important feature of this study is the use of an unsupervised chromatin-state representation rather than predefined chromatin-state annotations. Existing approaches such as ChromHMM and Segway have transformed genome annotation by assigning functional labels to genomic regions based on combinations of chromatin marks(Ernst and Kellis 2012; Hoffman et al. 2012). Our approach is complementary rather than competitive. Instead of optimizing genome segmentation, it represents each genomic interval by its integrated chromatin signature, allowing chromatin states to emerge directly from shared epigenomic similarity(Becht et al. 2018; McInnes 2018). This representation facilitates direct comparison across cell types, preserves relationships among genomic regions, and enables systematic identification of the chromatin features that contribute most strongly to epigenomic variation. We anticipate that this strategy will be particularly valuable for comparative epigenomic analyses in which identifying the major axes of chromatin-state remodeling is more important than assigning predefined functional annotations.

Our analyses further highlight practical considerations for integrating epigenomic and chromosome-conformation datasets. Because ChIP-seq and Hi-C interrogate genome organization at substantially different spatial resolutions, direct comparisons across genomic scales remain challenging. By examining the continuity of chromatin states, we found that extended genomic regions sharing identical chromatin signatures are relatively uncommon, explaining why integration was most informative for local architectural features such as chromatin loop anchors. Future studies combining higher-resolution chromosome-conformation data with expanded epigenomic profiles may enable similar analyses across larger structural domains, including compartments and chromosome territories.

Several limitations should be considered. First, the current study is based on two representative human cell lines and therefore does not establish whether the chromatin-state organization identified here is a general feature of tissues, developmental systems, or disease contexts. Expanding this analysis to additional cell types will be important for defining the generality of these organizational principles. Second, our conclusions are based on integrative analyses of existing epigenomic datasets and therefore do not establish causal relationships between individual chromatin features and chromatin-state organization. Functional perturbation of Polycomb complexes, CTCF, or individual histone modifications will be necessary to determine how these factors establish and maintain the chromatin states identified here. Finally, incorporation of additional epigenomic modalities, including DNA methylation, chromatin accessibility, transcriptional activity, and higher-resolution three-dimensional genome data, should provide a more comprehensive view of chromatin-state organization and its relationship to genome function.

In summary, our study demonstrates that integration of complementary epigenomic features resolves a small number of major chromatin-state classes that organize the human epigenome. These analyses reveal two complementary principles of chromatin organization: CTCF-associated chromatin defines a stable architectural framework for genome folding, whereas selective remodeling of Polycomb-associated chromatin constitutes the major source of cell-type-specific epigenomic divergence. Beyond establishing an analytical strategy for comparative epigenomics, this work provides a conceptual framework for understanding how architectural and regulatory chromatin states jointly shape genome organization across diverse biological contexts.

## Acknowledgements

We thank all the members of the Liu laboratory for discussion. We thank the assistance and support from the Biostatistics and Bioinformatics Facility at Fox Chase Cancer Center. Research reported in this publication was supported by the National Cancer Institute of the National Institutes of Health under Award Number P30CA006927. We acknowledge support from the National Institute of General Medical Science (R35GM154879 to Y.L.) and W. W. Smith Charitable Trust Grant (C2407 to Y.L.)

## Author Contribution Statement

Y.L. conceived and designed the project. J.M. and R.A.M performed all the analysis with supervision of Y.L. and Y.Z. Hi-C data was generated by Y.L. with assistance of R.G.S. Y.L. and Y.Z. supervised the project and Y. L. wrote the manuscript with J.M.

## Methods

Publicly available ChIP-seq datasets for HCT116 and K562 cells were obtained from the ENCODE Project. The following accession identifiers were used: for K562 cells, CTCF (ENCFF910QVN), H3K9me3 (ENCFF330EOT), H3K27me3 (ENCFF552AKP), H3K9Ac (ENCFF927XJD), and H3K27Ac (ENCFF488FYZ); for HCT116 cells, CTCF (ENCFF534GNR), H3K9me3 (ENCFF094ZHH), H3K27me3 (ENCFF977LFP), H3K9Ac (ENCFF960LSV), and H3K27Ac (ENCFF445BLD). Signal tracks were downloaded in bigWig format and all analyses were performed using the hg19 reference genome.

Genome-wide signal tracks were converted from bigWig to bedGraph format using the pybigwig Python library. Converted bedGraphs were partitioned into non-overlapping 1-kb genomic bins using the hg19 chromosome size annotations obtained from the UCSC Genome Browser. For each bin, signal intensity was calculated as the weighted average of all overlapping bedGraph intervals, using interval length as the weighting factor. Bins were indexed and mapped onto a genome-wide hg19 bin framework using pandas DataFrames, producing a unified matrix in which rows corresponded to 1-kb genomic bins and columns corresponded to chromatin binding profiles across both cell lines. Genomic bins lacking any signal across all datasets were excluded from downstream analysis.

Signal quality was assessed by generating histogram-based density plots of binned signal intensities for each chromatin feature. Density estimates were calculated using 45 bins to provide resolution across the full dynamic range of signal values. Log10-transformed density distributions were used to visualize low-frequency high-intensity values. Based on the characteristic three-phase distribution of signal across all datasets, bins with intensities below 2 were excluded as low-signal background, and high-intensity outliers were removed using a density threshold of fewer than 100 observations per bin. These thresholds were empirically determined and applied consistently across all datasets. Following filtering, each feature column was independently standardized using z-score normalization.

PCA was performed using the scikit-learn implementation in Python. The normalized feature matrix was transposed such that each chromatin feature in each cell line represented an observation across genomic bins. PCA was conducted using the first two principal components. Both genome-wide and chromosome-level analyses were performed; for chromosome-level PCA, the normalized matrix was subset by chromosome prior to analysis. Pairwise Euclidean distances in PC space between corresponding chromatin features in HCT116 and K562 cells were calculated to quantify epigenomic divergence at the whole-genome and chromosome-specific levels. Chromosome-level distance values were compiled into a feature-by-chromosome matrix and visualized as hierarchically clustered heatmaps generated in R. UMAP was applied to the normalized genomic bin matrix using the umap-learn Python package. Embeddings were generated using 100 nearest neighbors, a minimum distance of 0.01, and Euclidean distance as the metric. Both whole-genome and chromosome-specific embeddings were generated. Individual chromatin feature signal intensities were overlaid onto UMAP embeddings as continuous color scales using the colorcode function, enabling identification of feature-specific enrichment patterns within and across clusters. Cluster-level chromatin signatures were summarized using heatmap representations in which relative signal intensities for each feature were compiled per cluster.

### Hi-C experiments and data analysis

Both K652 and HCT-116 cells were purchased from ATCC. K562 and HCT-116 cells were cultured in 10cm dishes with RIPM medium and McCoy’s 5A medium, respectively, at 37°C in 5% CO2. Both mediums contains GlutaMAX supplement (Gibco, 36600021) supplemented with 10% FBS (Gibco, 16000044) and 1% penicillin-streptomycin (Gibco, 15140). DMSO (#D2650) was purchased from Sigma-Aldrich, USA and the cells were treated with 10ul DMSO for three hours followed by formaldehyde fixation.

Hi-C for fixed cells was performed as described in our previous study(Liu and Dekker 2022). Briefly, the fixed cells were first lysed to obtain nuclei. After being washed twice with cold NEBuffer 3.1, the nuclei from fixed cells were resuspended in 342ul NEBuffer 3.1 with 0.1% SDS and the tube was gently mixed. The tube was then incubated at 65°C for 10min and put on ice immediately, followed by the addition of 43ul of 10% Triton X-100 and gentle mixing. DpnII (400U) digestion was performed at 37°C overnight with gentle rocking. Once enzyme digestion was completed, the reaction was incubated at 65°C for 15mins to inactivate DpnII. DNA overhanging ends were then filled in with biotin-14-dATP at 23□°C for 4□hours and then ligated with T4 DNA ligase at 16□°C for 4□hours. DNA was treated with proteinase K at 65□°C overnight to remove proteins. Ligation products were purified, fragmented by sonication to an average size of ~200□bp and size-selected to fragments of 100–350□bp. We then selectively purified biotin-tagged DNA using streptavidin beads before performing end repair, dA-tailing and adaptor addition, using NEBNext Ultra II DNA Library Prep Kit for Illumina (E7645L). Dual indexes were then added by PCR using NEBNext Multiplex Oligos for Illumina (E7600S). The PCR primers were removed from final libraries using AMPure beads. Hi-C libraries were then sequenced using PE150 bases on an Illumina HiSeq 4000 or an Illumina NovaSeq instrument.

Sequencing data was first trimmed to PE50 using the in-house scripts. All Hi-C PE50 fastq raw sequencing files were mapped onto hg19 human reference genome using distiller-nf mapping pipeline (https://github.com/mirnylab/distiller-nf). After mapping, aligned reads were further processed to remove duplicates (https://github.com/mirnylab/pairtools) to obtain a set of filtered reads defined as valid pairs. Valid pairs were then binned into contact matrices at 20□kb and 100□kb resolutions using cooler50. Intrinsic Hi-C biases were removed using the Iterative Correction and Eigenvector decomposition (ICE) procedure(Imakaev et al. 2012). This was applied to all of the matrices, ignoring the first two diagonals to avoid short-range ligation artifacts at a given resolution, and bins with low coverage were removed using the MADmax filter with default parameters. Contact matrices were stored in ‘.mcool’ files and used in downstream analyses.

Anchors of CTCF-CTCF loops were previously identified from both cell lines (Sanborn et al. 2015; Rao et al. 2017). In total, 3169 and 6058 loops were identified from HCT-116 and K562 cells(Rao et al. 2014; Rao et al. 2017). The center of loop anchor were used to obtain 1-kb loop anchor bins. The loop anchor bins were then mapped on the five clusters.

To identify TAD boundaries, we first calculated observed/expected Hi-C matrices of each sample for 20 kb binned data, correcting for average distance decay as observed in the *P*(s) plots (cooltools compute-expected). We then aggregated the observed/expected Hi-C matrices of each sample at the TAD boundaries that were identified from the sample without any treatments, covering 600kb up and downstream of each boundary, and then generated a pileup heatmap of TAD boundaries for each sample.

All Python scripts used for bigWig conversion, genome binning, quality control, normalization, PCA, and UMAP analysis, as well as the R script used for heatmap generation, are available upon reasonable request from the corresponding author.

